# The Tell-Tale Genome

**DOI:** 10.1101/091074

**Authors:** Eugenio Bortolini, Luca Pagani, Enrico R. Crema, Stefania Sarno, Chiara Barbieri, Alessio Boattini, Marco Sazzini, Sara Graça da Silva, Gessica Martini, Mait Metspalu, Davide Pettener, Donata Luiselli, Jamshid J. Tehrani

## Abstract

Observable patterns of cultural variation are consistently intertwined with demic movements, cultural diffusion, and adaptation to different ecological contexts (Cavalli-Sforza and Feldman 1981; Boyd and Richerson 1985). The quantitative study of gene-culture co-evolution has focused in particular on the mechanisms responsible for change in frequency and attributes of cultural traits, on the spread of cultural information through demic and cultural diffusion, and on detecting relationships between genetic and cultural lineages. Here, for the first time, we make use of worldwide whole-genome sequences (Pagani et al. 2016) to assess the impact of demic diffusion on cultural diversity, focusing on the variability observed in folktale traditions (N=596) (Uther 2004) in Eurasia and Africa. We show that at small geographic scales (<=5000 km) there is a strong correlation between folktale and genomic distance when the effect of geography is corrected, while geographic distance has no independent effect on the distribution of folkloric narratives at the same spatial scale. This points to demic processes (i.e. population movement and replacement) as the main driver of folktale transmission at limited geographic ranges. The role of population movements becomes more apparent when regions characterized by episodes of directional expansions, such as the Neolithization of West Eurasia, are examined. Furthermore, we identify 89 individual tales which are likely to be predominantly transmitted through demic diffusion, and locate putative focal areas for a subset of them.

## 1 Introduction

Advances in DNA sequencing have opened new ways for exploring the demographic histories of human populations and the relationship between patterns of genetic and cultural diversity around the world. Newly available genome-wide evidence goes beyond the use of linguistic relationship as a measure of common ancestry (Currie *et al.* 2010, Ross *et al.* 2013, Mathew and Perreault 2015, da Silva and Tehrani 2016), and offers unprecedented support for studying the mechanisms underlying the transmission of cultural information over space and time (Cavalli-Sforza and Feldman 1981, Boyd and Richerson 1985, Collard *et al.* 2006, Ackland *et al.* 2007, Pinhasi and von Cramon-Taubadel 2009, Gray *et al.* 2010, Fort 2012, Lycett 2015), as well as the coevolution of genetic and cultural traits (Ammerman and Cavalli-Sforza 1984, Renfrew 1992; 2001, Bell *et al.* 2009, Itan *et al.* 2009, Creanza *et al.* 2015, Haak *et al.* 2015) across populations.

A key question for research in this area concerns the extent to which patterns of cultural diversity documented in the archaeological and ethnographic records have been generated by demic processes (i.e. the movement of people carrying their own cultural traditions with them) or cultural diffusion (i.e. the transfer of information without or with limited population movement/replacement) (Collard *et al.* 2006, Crema *et al.* 2014, Fort 2015). Demic and cultural diffusion are not mutually exclusive conditions, rather opposite extremes of a continuous gradient whose intermediate and composite positions more accurately represent empirical reality.

A broadly adopted null model against which instances of demic processes have been assessed draws on the expectation that selectively-neutral cultural or genetic variants would form geographic clines produced over time by Isolation-by-Distance processes (IBD; (Wright 1943)). Under an IBD model individuals or groups which are spatially closer to each other are expected to be more similar than individuals or groups that are located further apart. A positive correlation between cultural or genetic dissimilarity and geographic distances between samples is therefore used to infer processes of cultural transmission of non-adaptive information without population replacement (Pinhasi and von Cramon-Taubadel 2009). However, observed genetic distance is the composite result of serial founder events (SFE), long term IBD and subsequent migratory events, which imply recent movement and resettling of people. The latter component is therefore the one that can be used to infer the relative effect of demic processes when looking at the distribution of cultural variants that may have spread over the past few thousand years.

In a recent study Creanza and colleagues (Creanza *et al.* 2015) investigated the process responsible for the observed global distribution of (phonetic) linguistic variability by comparing it to genetic and geographic distances. The authors found high correlation between genetic and geographic distances at a worldwide scale, while linguistic distances were spatially autocorrelated only within a range of ~10000 km. The lack of residual correlation between genetic and linguistic distances up to this spatial scale was interpreted as a signal of cultural diffusion being the main driver of the distributions of phonetic variants in human populations.

The use of genetic variability to infer the relative impact of demic processes on the distribution of cultural traits therefore depends on being able to disentangle genetic signals from geography. The high correlation between genetic and geographic distances at a global scale (Ramachandran *et al.* 2005) lowers the inferential power of this model. However, this relationship is not constant across different geographic scales. The correlation obtained using both pairwise geographical distances and genetic split times is stronger when measured across all possible population pairs at larger geographic scales, while it is considerably lower at smaller geographic distances (below ~8000 km for the present dataset), possibly because of more recent and short-range population movements (Figure 1 top). It is worth noting that global trends have been forming over the past ~40000 years, while most cultural traditions are likely to have evolved more recently. This is supported by previous studies (Creanza *et al.* 2015), and suggests that the effect of population movements independent from IBD can be identified only within limited geographic scales (<8000 km). At this spatial resolution events shaping the distributions of genetic and cultural divergence are more likely to occur at the same temporal scale and, hence, to be more probably causally related.

**Figure 1:**
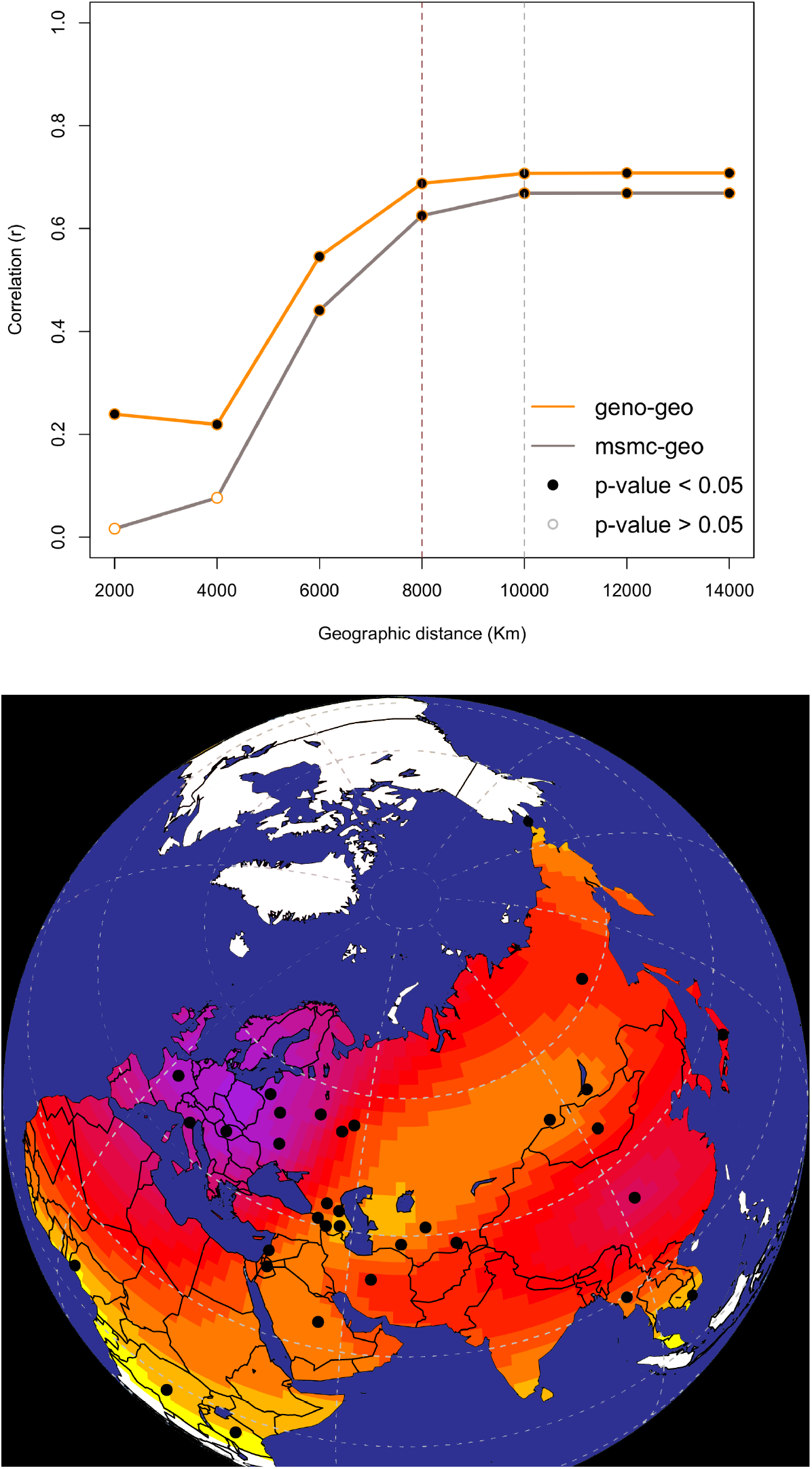
Top) Plot of linear correlation values between pairwise genetic distance (based on whole genome) and pairwise geographic distance, and between genetic split-times (MSMC) and geography over cumulative bins of geographic distance; Bottom) Map showing the spatial distribution of 33 populations comprised in Dataset_MAIN_. Surface colours represent interpolated richness values (i.e. the number of folktales exhibited by each population). Purple indicates higher values, while yellow indicates coordinates with the lowest number of folktales (in the present sample). The distribution of richness colours clearly highlights the fact that currently available data on folkloric traditions are skewed in favour of European and east-Asian lineages.

Here we capitalize on the short-range decoupling of genetic and geographic distance to further infer mechanisms of genetic and cultural coevolution by using newly available genomic evidence (Pagani and et al. in press) as an unbiased proxy of population movement. To do so, we investigate the relative impact of demic processes and cultural diffusion on the observed distribution of a set of individual folktales in Eurasia and Africa. Folktales are a ubiquitous and rigorously typed form of human cultural expression, and hence particularly well-suited for investigating cultural processes at wider cross-continental scale. Researchers since the Brothers Grimm (Grimm 1884) have long theorized about possible links between the spread of traditional narratives and population dispersals and structure, but have found mixed levels of support for this hypothesis when using indirect evidence for demic processes, such as linguistic relationships among cultures. One recent study suggested that the distributions of a substantial number of fairy tales shared among Indo-European populations were more consistent with linguistic relationships than their geographical proximity, suggesting they were inherited from common ancestral populations (da Silva and Tehrani 2016). Another study found evidence of both demic and cultural diffusion in the distributions of folktales recorded among Arctic hunter-gatherers (Ross and Atkinson 2016), while a third suggested that clinal patterns are better explained by geographic distance than linguistic barriers in variants of the tale of The Kind and Unkind Girl in European groups (Ross *et al.* 2013).

In the present study we focus on 596 folktales comprising “Animal Tales” and “Tales of Magic” (Uther 2004), typed as present (1) or absent (0) in 33 populations (Dataset_MAIN_) for which whole-genome sequences are available and exhibiting presence of at least five folktales (Figure 1 b; SI1 Section 1). We hypothesize that if the extent of folktale sharing between populations positively correlates with genomic similarity, the diffusion of folkloric traditions can be predominantly explained by population movements and admixture between demes (demic processes). At the same time, the decline of similarity in folktale traditions between populations at increasing geographic distance (IBD) needs to be contextualized in light of human interaction and cultural transmission without population replacement.

We test our hypothesis by using both a frequentist hypothesis-testing approach and a model-selection approach to ascertain the proportion of folktale variability that is explained by genetic distance as well as the proportion that is more significantly dependent on geographic distance. We further check our results in a controlled case study of small-scale demic diffusion by looking at a subset of populations affected by Neolithic and post-Neolithic migrations from western Asia to Europe. In addition, we attempt to identify individual folktales likely to be predominantly transmitted through demic processes, and try to suggest their potential focal area intended as a putative proxy for their center of origin.

## 2 Results

### 2.1 Competing effects of genetic and geographic distance on folktale variability

A Model Selection approach based on Akaike’s Information Criterion (AIC, (Akaike 1973, Burnham and Anderson 2002); SI1 Section 3) confirms the impossibility of disentangling the effect of geography over genetic distance to predict sharing of folktales between population pairs at a global scale (Table 1; SI1 Section 4) by identifying the combined model comprising both variables as the best one. The same is true when focusing on Eurasian populations alone (Dataset_Eurasia_, N=30) to control for the effect of the Out of Africa expansion on genetic distance. However, when considered separately, AIC Δs (SI1 Section 3) indicate that genetic distance outperforms geographic distance, with both Akaike weights and likelihoods confirming that genomic variability has a higher goodness-of-fit than geography alone. When examined at different cumulative spatial scales, genomic distance has a predominant role (over both the combined model and geography alone) at the smallest spatial scale (< 2000 km) as well as when all population pairs within a distance of 10000 km are considered. For all remaining distance bins the combined model provides the best explanation, while geography alone has the lowest goodness-of-fit (Table 1).

**Table 1:**
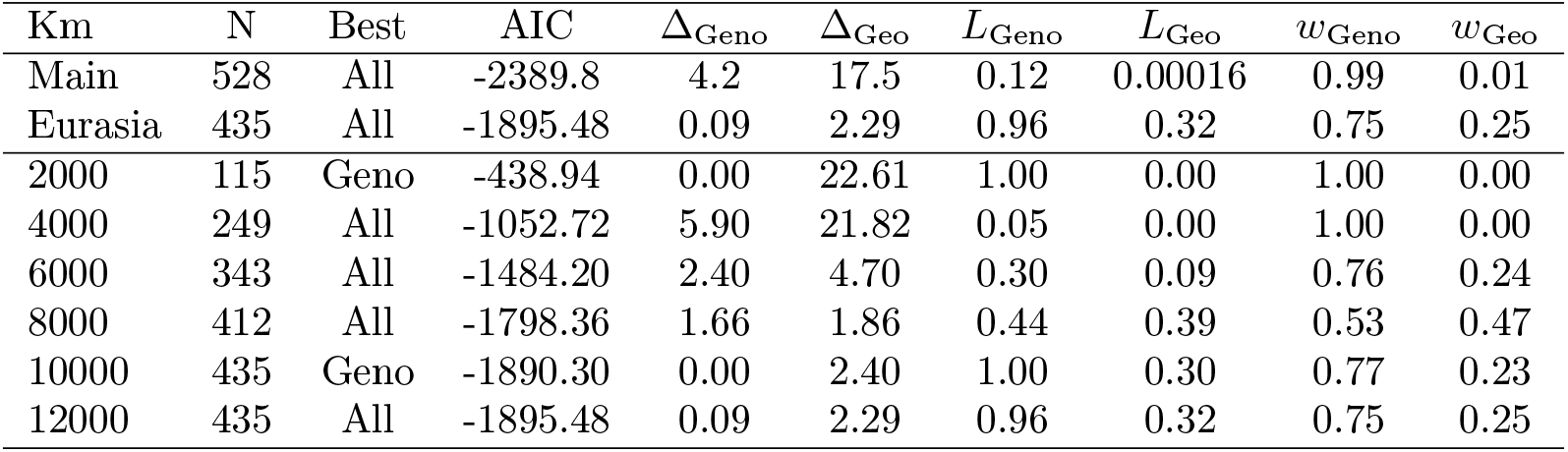
Model selection at cumulative geographic distances

### 2.2 Genetic-folktale relationship at different geographic scales

The predictive power of genetic distance on the folktale diversity is further investigated in Dataset_EURASIA_ over cumulative geographic distance through linear correlation. Correlation values are high and significant up to ~6000 km (Figure 2a, red line) and rapidly decrease towards global values with increasing geographic distance. Conversely, the correlation between folktales and geography is lower and nearly constant at any bin (Figure 2a, blue line). As predicted by preliminary results (Figure 1, top), moderate geographic distances allow us for a more efficient disentanglement between genetic and geographic distance, suggesting that at this scale the strong relationship between genetic variability and folk narratives is likely to be independent of the underlying spatial distribution. When looking at the same pattern for individual geographic bins via Multivariate Partial Mantel Correlograms (Figure 2b), residual significant correlation can only be observed between folktale and genomic distance for bins up to 4000 km, providing a lower geographic boundary for the inferred process of demic diffusion.

**Figure 2:**
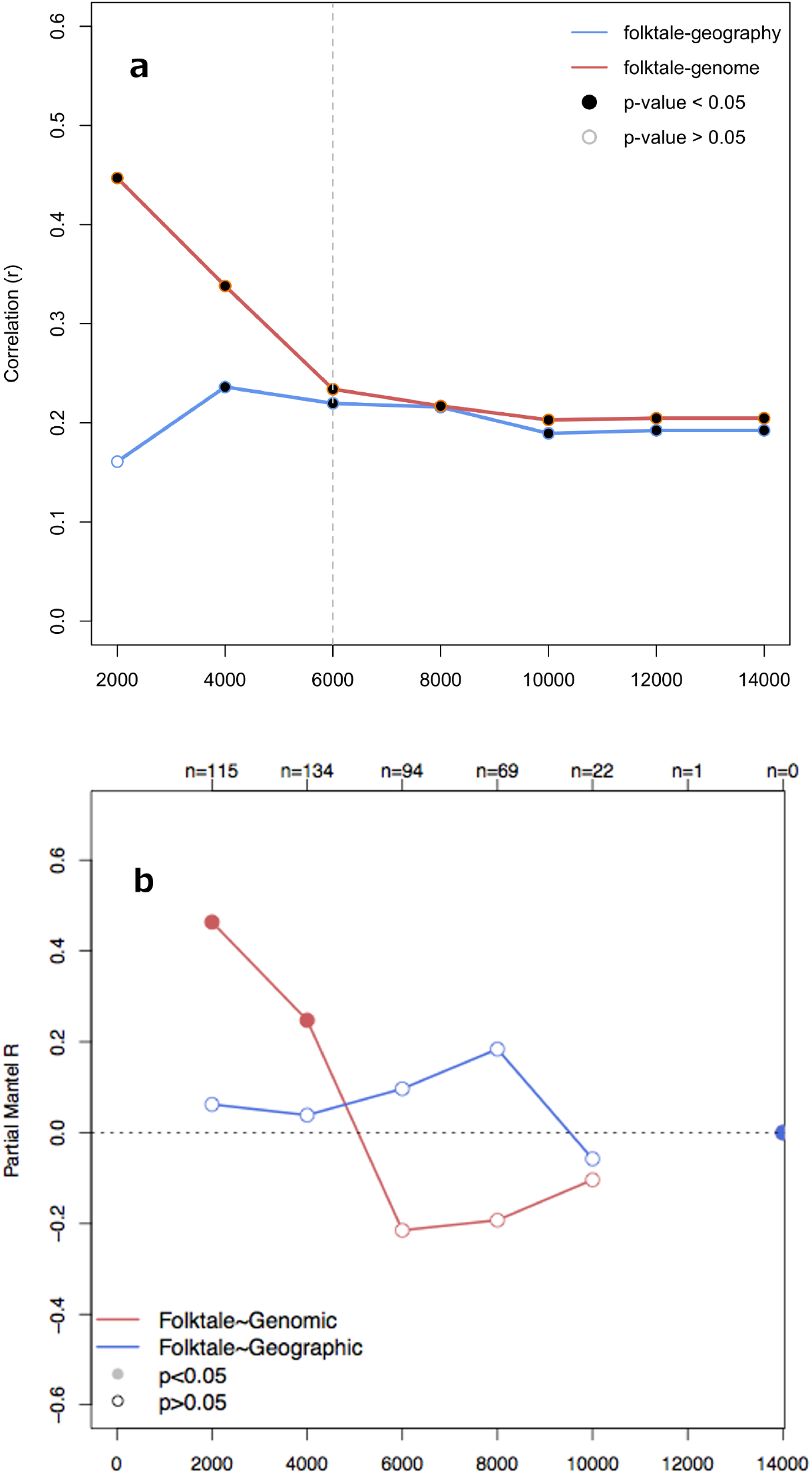
a) Linear correlation between folktale distance and genetic distances based on whole genome and allele frequency intervals, and between folktale distance and geographic distance. Correlation values are calculated over cumulative geographic distances comprised between 2000 and 14000 km; b) Multivariate Partial Mantel Correlogram showing the residual linear correlation between folktale distance and genetic distances after controlling for geographic distance, and between folktale distance and geographic distance after controlling for genetic distance. Correlation values are calculated for individual geographic bins.

### 2.3 Case study: Neolithic and post-Neolithic expansions in West Eurasia

We hypothesize that the observed scale-dependent fit between genes and folktales may be the result of recent demic movements. To test this hypothesis, we focus on a subset of samples (N=8; Dataset_EURASIA_POSTNEO_; SI1 Section 1) which was likely affected by the Neolithic/post-Neolithic expansions in Europe, one of the best documented cases of recent demic diffusion (Ammerman and Cavalli-Sforza 1984, Gkiasta *et al.* 2003, Pinhasi *et al.* 2005, Fort 2012; 2015). Correlation between folktale and genetic distance is considerably higher than when the whole of Eurasia is considered (r=0.85, p<0.0001) and robust to permutation tests (SI 1 Section 5, Figure S1-5.1). This suggests that traditional narratives, as much as other cultural and technological innovations, may have spread by means of demic diffusion during Neolithization (da Silva and Tehrani 2016). Phylogenetic (Neighbor-Joining) and NeighborNet analysis of folktale distance on the same subset of populations cement this inference. The clear tree-like structure of the latter (*δ* score=0.18, Q-residual=0.003; (Holland *et al.* 2002); Figure 3) is comparable to results obtained for other cultural and linguistic datasets (Tehrani 2013) and is consistent with a scenario of dispersal of modern folklore in western Eurasia as the result of branching processes, with appreciable reticulation only within linguistic sub-families. This is consistent with recent evidence of post-Neolithic demic movements being a putative vector of present-day linguistic variability in Europe (Haak *et al.* 2015, Allentoft *et al.* 2015).

**Figure 3:**
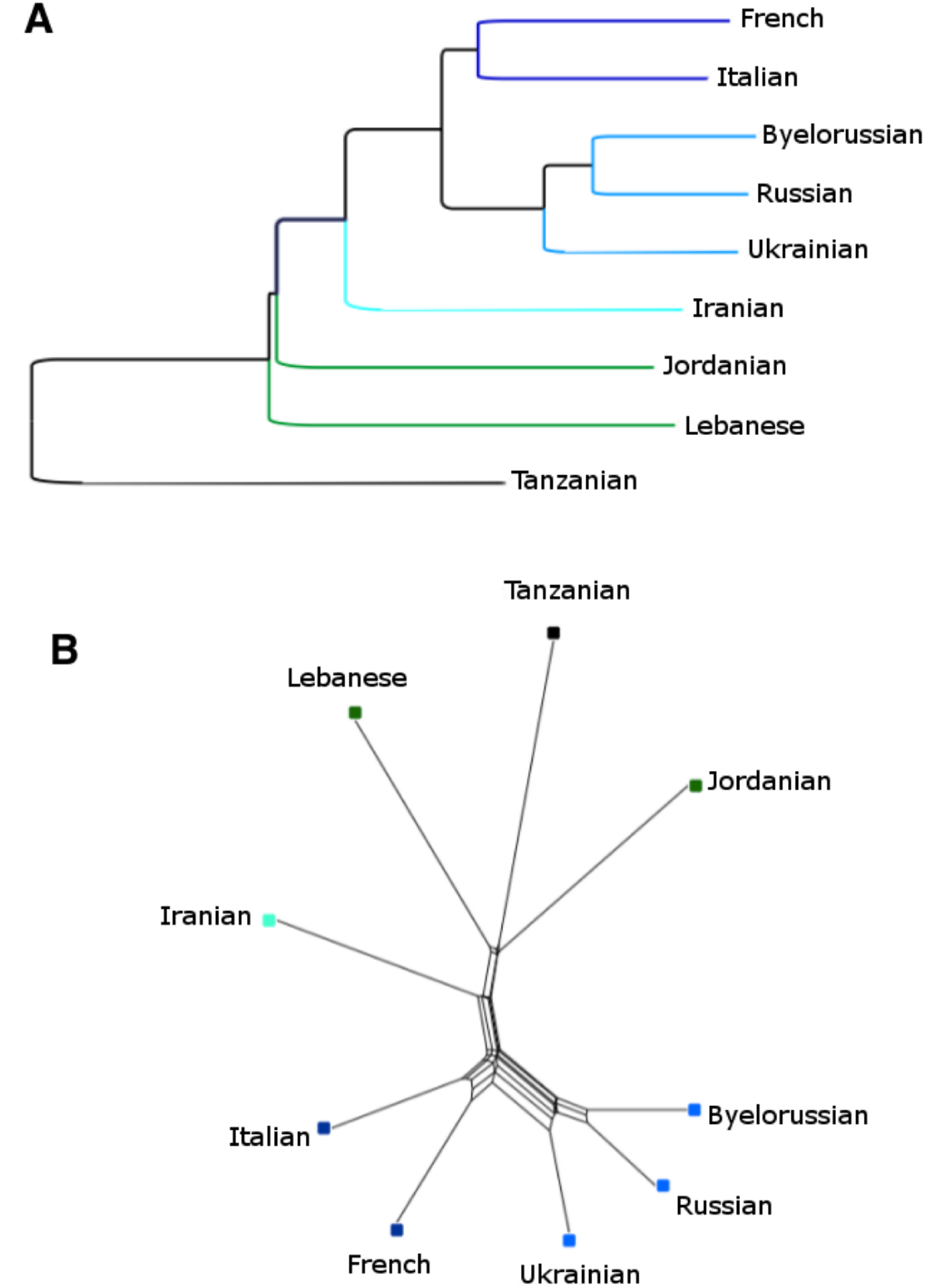
A) NJ tree calculated on folktale distances comprising Neolithic/post-Neolithic populations and Tanzania as the outgroup. Colours identify linguistic families; B) NeighborNet calculated on the same subset of populations.

### 2.4 Transmission of individual folktales

When a given folktale is transmitted through population movement and replacement, all the populations that share it should exhibit significantly lower genetic distance than populations that do not present with that specific tale. Following this assumption, we search for folktales that are more likely to be transmitted via demic processes. We achieve this by comparing the distribution of pairwise genetic distances among populations sharing a given tale against the distances of the remaining pairs of populations using a Mann-Whitney-Wilcoxon test. We focus on the 308 folktales that are present in at least five populations and run two separate tests, the first considering all pairs of populations, and a second considering only those within a geographic range of 5000 km — the mid-point between the threshold distances of 4000 and 6000 km identified earlier. Of the 308 analyzed folktales, 89 (30%) present with significantly lower than expected pairwise genetic distance. Among these 89 tales, 63 are more likely to be driven by population movements at small geographic scales (i.e. the signal is only significant below 5000 km), while it is impossible to determine the most probable transmission mechanism at longer distances. Of the remaining twenty-six folktales, twenty-two have been probably traveling with human groups within a maximum range of 5000 km, while the last four tales are possibly prone to demic diffusion both below and beyond 5000 km.

### 2.5 Folktale dispersal and focal areas

For a subset of the analyzed folktales we identify focal areas, representing potential areas of origin and defined as locations that maximize the decay of a given folktale abundance over geographic distance measured with Pearson correlation coefficient (SI1 Section 6). Focal areas were generated for the four folktales which are likely to have been predominantly transmitted through demic processes at a global scale, and for the 16 most widespread folktales that show no evidence of demic diffusion (SI1 Section 6). As far as the four “demic” tales identified above (ATU4, ATU248, ATU280A, and ATU650A), three present a consistent pattern in which eastern Europe/Caucasus is the most probable focal area, whilst the fourth (ATU248) had most likely its point of origin in northern Asia (Figure 4).

**Figure 4:**
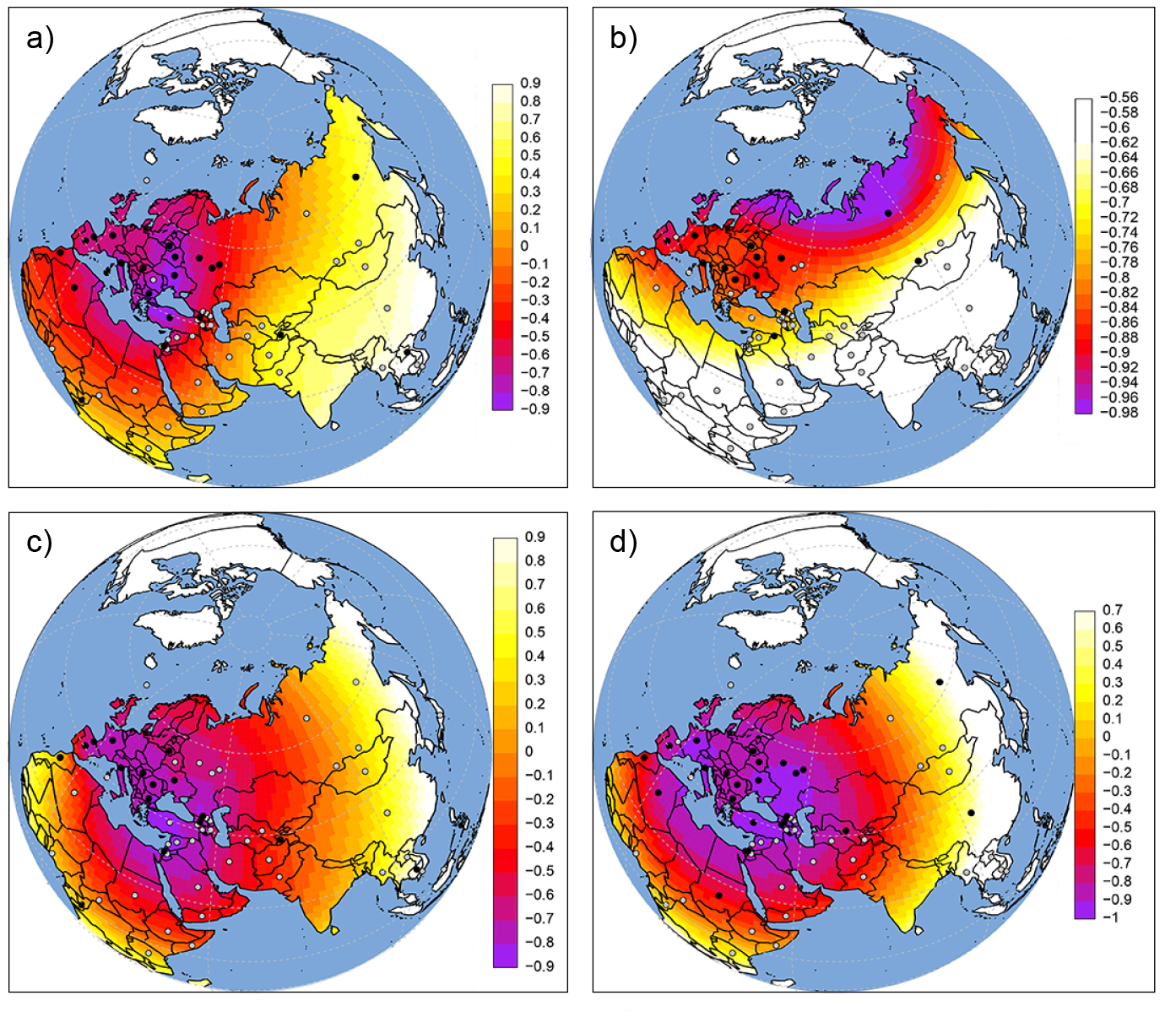
Four examples of plausible origin and dispersion patterns for tales likely to have been predominantly spread through demic diffusion: a) ATU4; b) ATU248; c) ATU280A; d) ATU650A.

The most widespread or “popular” folktales which do not show any evidence of demic transmission can be grouped into four main focal areas. Some of these tales possibly started to be diffused mostly via cultural transmission from eastern Europe (such as ATU155, The Ungrateful Snake Returned to Captivity; Figure S1-6-I a), while others are more probably starting their journey from Caucasus (Figure S1-6-I f). Examples of the latter are ATU400 “The Man on a Quest for His Lost Wife”, ATU480 “The Kind and Unkind Girls”, ATU531 “The Clever Horse”, and ATU560 “The Magic Ring”. Some narrative plots might have been originated in northern Asia - such as the famous “Thumbling” (Tom Thumb; Figure S1-6-I o), while a last group could have spread from Africa (Figure S1-6-I n), as in the case of ATU670 “The Man Who Understands Animal Language”.

## 3 Discussion

Folktales are universal cultural expression linked to various vectors of propagation over generations and across geographic barriers, that allows us to address questions on cultural evolutionary processes at a cross-cultural and cross-continental scale. Our results provide new insights on the processes driving the spread of folkloric narratives by using newly generated genomic evidence as a proxy of population movements that goes beyond previous studies that were limited to a single language family (da Silva and Tehrani 2016).

Distance between populations based on folktales significantly correlates with genomic distance at a cross-continental level, suggesting an important role for population movements in the spreading of these tales. However, at broad geographic scales it is impossible to effectively disentangle the effect of geographic distance over genetic distance, and to use these measures to estimate the different contribution of demic and cultural variability.

More definitive results are in fact obtained at small geographic scales (below 4000-6000 km), where folktale distance strongly correlates with genetic distance even after controlling for geography, suggesting the dominance of demic processes (population movement and replacement) within a radius of 5000 km. Beyond this threshold a major impact of demic processes can only be inferred for a very limited number of tales (ATU4, ATU248, ATU280A, and ATU650A; Figure 4). Model selection definitively shows that genetic distance provides a better fit than geographic distance and indicates that the demic component of folktale transmission is up to three times more likely than cultural transmission at most geographic scales, even where we find only little evidence of residual correlation due to the noisy nature of cultural evidence. While model likelihood and weights make it clear that folktale transmission is always the result of co- occurring mechanisms in different relative proportions, they suggest that demic processes are predominant below 5000 km, where transmission may be independent from the underlying spatial structure in the dataset. Notably, such a geographic range is shorter than the maximum predicted by our (Figure 1a) and previous results (Creanza *et al.* 2015), hence showing that the observed pattern is not due to saturation in the available data.

When tested in a controlled case study of well documented and small-scale human migrations (such as Neolithic and post- Neolithic expansions into West Eurasia) results are further supported by higher correlations and evidence of branching among the interested demes, suggesting that folktales may have been part of the many cultural packages adopted and elaborated in this context, and supporting a view of the Neolithization of Europe as a composite process predominantly driven by demic diffusion and continuous dispersal from Western Asia and eastern Europe (Pinhasi and von Cramon-Taubadel 2009, Fort 2015, Hofmanova *et al.* 2016). The long-range patterns detected by our analyses (Figure 4; Figure S1-6-I) may complement this picture by suggesting a more ancient origin of some of these folktales (SI1 Section 6; (Bottigheimer 2009; 2014, Thompson 1977, Propp 1968)). On a broader level, these trends can be used in the future to infer directional trends of folktale diffusion, as well as to test for the emergence of systematic social biases or cultural barriers whose chronology may be independently ascertained.

Our results also have a wider resonance with broader questions concerning cultural evolution and the mechanisms of cultural transmission over time and space. In particular, they show that the use of newly generated, whole-genome sequences offers a unique opportunity for an unbiased assessment of the temporal scale and relevance of demic diffusion events to patterns of cultural variation in the ethnographic and archaeological records. Nevertheless, they also offer a cautionary tale concerning the investigation of such events at a cross-continental or global scale, where geographic clines in genetic variability are the result of different processes that can hardly be disentangled and that may present with considerable temporal mismatch with more recent cultural processes.

## Materials and Methods

### Dataset description

Folktale data were sourced from the Aarne Thompson Uther catalogue (ATU; (Uther 2004)). The present dataset comprises Animal Tales (ATU 1-299), and Tales of Magic (ATU 300-749). Of the 198 societies in which the tales were recorded, 73 matched available genetic data. Of these, 33 populations exhibiting at least 5 folktales were selected (Table S1-1, Figure 1b). Each population is described by a string listing the presence (1) or absence (0) of any of the included 596 folktales (Table S1-2).

### Genetic distance

Genetic distances were estimated by the average pairwise distances between two genomes, one from each population, including both coding and non-coding regions to avoid ascertainment biases. Genetic distance for (i, j) pairs of populations represented by more than one genome was calculated as the average of all possible (i, j) pairs of genomes. As a consequence the diagonal of the genetic distance matrix was not constrained to be zero.

### Multiple sequential Markovian coalescent model analysis

Multiple sequential Markovian coalescent model (MSMC; (Schiffels and Durbin 2014) was used to infer genetic split times between the 528 pairs of populations comprised in the original dataset (Pagani and et al. in press), taking one genome each as representative. Genetic split times were inferred at a cross coalescent rate of 0.5 and compared with folktale and geographic distances as an additional measure of genetic similarity.

### Folktale distance

Folktale distance between population pairs was calculated as asymmetric Jaccard distance (Jaccard 1901).

### Geographic distance

Geographic distance was calculated as pair- wise great circle distance with a waypoint located in the Sinai Peninsula to constrain movement of African demes (through the package gdistance in R; (van Etten 2014)). Coordinates (longitude and latitude in decimal degrees) identify the assumed center of the area occupied by a given folkloric tradition as defined by the ATU index.

### Measures of statistical association

The relationship between folktale, genetic, and geographic pairwise distance matrices at cumulative geographic scales was assessed through linear correlation analysis. Multivariate Partial Mantel Correlograms were used to measure and plot residual correlation between folktale and genetic/geographic distance after controlling for the remaining variable in individual geographic bins. All the analyses were performed using the ecodist package in R (Goslee and Urban 2007).

### Model selection

Model selection was based on Akaike’s Information Criterion (AIC; (Akaike 1973, Burnham and Anderson 2004); SI1 Section 4) calculated via linear regression analysis with stepwise variable selection using the stepAIC function of the package MASS in R (Venables and Ripley 2002). Model weights were calculated using the package MuMlN in R (Barton 2016). Models comprised: i) modelALL, a composite model consisting of genetic distance + geographic distance; ii) modelGeneticsOnly; iii) modelGeographyOnly.

### Mechanisms acting on individual folktales

Mann-Whitney-Wilcoxon test was used to test for populations sharing a specific tale having significantly lower genetic distance than populations not exhibiting the same tale. Cases which were significant after Bonferroni correction were considered for inferring processes of transmission. Only folktales that were present in at least five populations were considered. The analysis was performed for a buffer of 5000 km and at a global scale (15000 km).

### Folktale dispersal

Population pairs were binned into fixed intervals of geographic distance (2000 Km), and for each bin the proportion of populations exhibiting a given folktale was calculated. The obtained percentage was then linearly correlated with geographic distance from all the populations exhibiting the same folktale. The surface distribution of correlation coefficients was obtained through plate spline interpolation using the fields package (Nychka *et al.* 2016) in R (Team 2016).

## Acknowledgments

The authors are grateful to Adrian Timpson, Anne Kandler, Dugald Foster, Jeremy Kendal, and Rachel Kendal for their comments and useful suggestions. EB is supported by SimulPast Consolider Ingenio project (CSD2010-00034) funded by the Spanish Ministry of Science and Innovation.

